# Barley (*Hordeum vulgare*) maintains tricarboxylic acid cycle activity without invoking the GABA shunt under salt stress

**DOI:** 10.1101/2025.11.06.687118

**Authors:** Ali Bandehagh, Nicolas L. Taylor

## Abstract

Plants maintain energy balance under salinity stress through increased respiration and energy use, processes also associated with reactive oxygen species generation. Although respiration imposes a high energy cost, mitochondrial respiration and the tricarboxylic acid (TCA) cycle activity are vital for ATP production and providing electron donors that drive ion exclusion and ROS detoxification. This study examined the molecular basis of respiratory responses to salinity in barley using physiological, biochemical, metabolomic, and proteomic analyses. Salt exposure resulted in sodium accumulation, decreased photosynthesis and biomass, and increased respiration. Metabolite profiling indicated activation of the TCA cycle, while proteomics showed increased abundance of all targeted TCA enzymes, including phosphoenolpyruvate carboxylase isoforms and succinate dehydrogenase. Enhanced pyruvate oxidation and accumulation of downstream metabolites suggested that the classical TCA cycle underpins barley’s salinity tolerance. Conversely, reduced levels of 2-oxoglutarate and succinate implied limited activity of the GABA shunt. The absence of detectable arginine and ornithine, unlike their salt-induced increase in wheat, further indicated that the GABA shunt contributes minimally to barley’s salinity tolerance. Overall, barley primarily relies on enhanced mitochondrial respiration and the canonical TCA cycle, rather than GABA shunt metabolism, to cope with salinity stress.

**Highlight:** Barley sustains respiration via the TCA cycle rather than activating the GABA shunt under salt stress, whereas in wheat, salt physiochemically inhibits TCA cycle function, and the GABA pathway is induced.

**Graphical Abstract:** 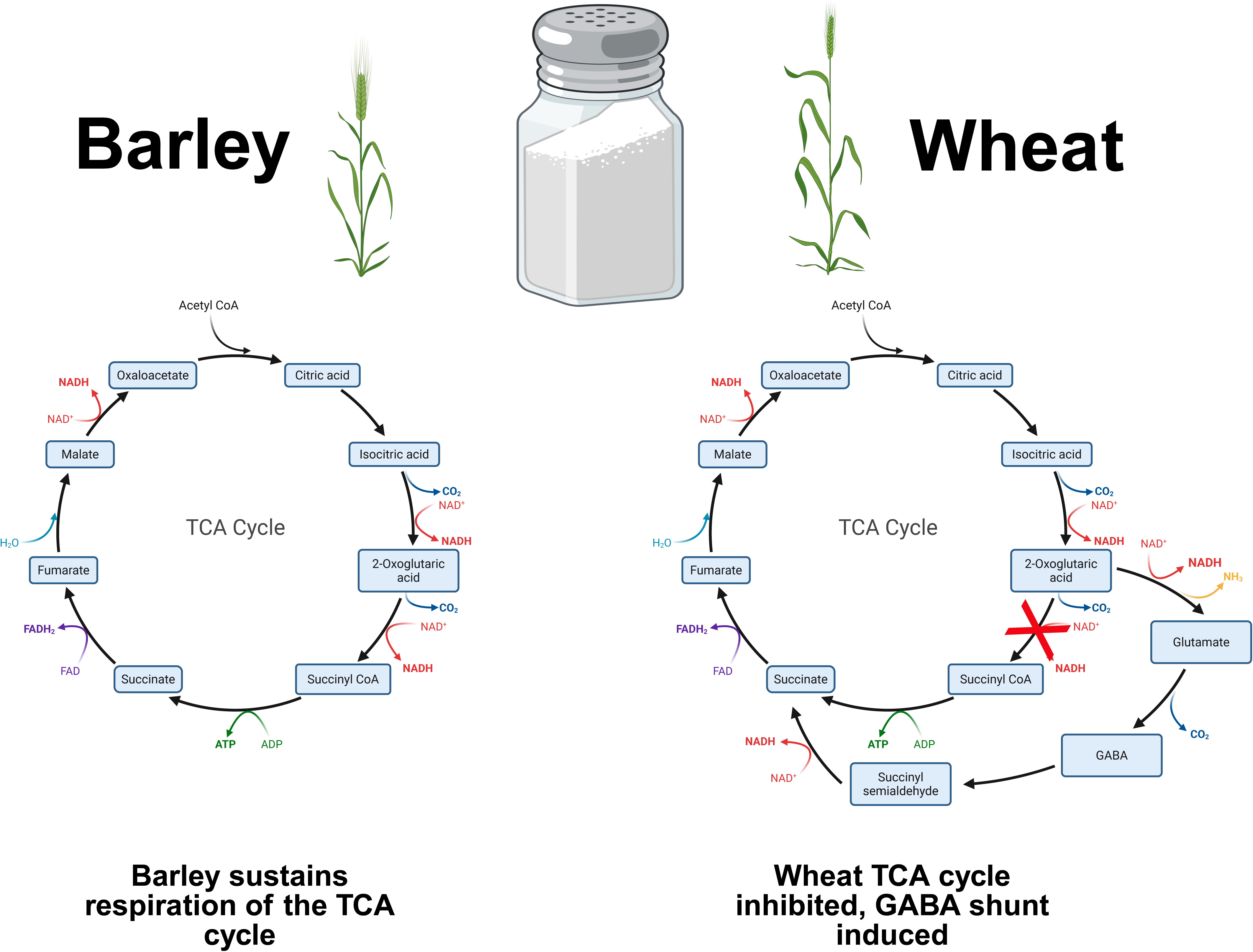

## Introduction

Salinisation of soils poses a significant challenge for crop production, and the area of salt-affected soils continues to grow, especially in irrigated lands (Machado and Serralheiro, 2017). Most energy captured during photosynthesis is stored in carbon compounds used for plant maintenance (Amthor, 2000; Jacoby *et al*., 2011). Salt exposure hampers growth, limits photosynthesis, and redirects respiration towards defence. Plants adopt various strategies to regulate carbon balance and energy distribution under salt stress, resulting in tissue and osmotic tolerance. Glycolysis in the cytoplasm, the TCA cycle, and the electron transport chain in mitochondria utilise energy stored in photosynthetic sugars to produce ATP, which fuels many cellular biochemical reactions (Tiwari *et al*., 2002). Salt exposure increases ATP demand because ATP is essential for several energy-intensive adaptive processes, including ion homeostasis, ROS protection, and osmotic regulation. During salinity stress, plants tend to accumulate sucrose and amino acids, especially proline, alanine, GABA, and lysine, alongside notable degradation of organic acids, particularly TCA intermediates in wheat (Che-Othman *et al*., 2019), barley (Widodo *et al*., 2009), and other crops (Sanchez *et al*., 2008).

Plants use different strategies to enhance stress tolerance by maintaining energy homeostasis under salinity. Many of these strategies demand increased energy use and respiration, which are vital for plant survival. A plant’s energy balance is determined as the photosynthesis-fixed carbon minus respiratory carbon, with the net assimilated carbon supporting growth. Under saline conditions, respiration can slow, and the plant’s net carbon status diminishes due to a significant drop in photosynthesis (Schwarz and Gale, 1981). Salt stress reduces chlorophyll levels, leading to decreased photosynthesis; however, experiments indicate that specific respiration rates can increase (Che-Othman *et al*., 2019; Reuveni *et al*., 1997; Schwarz and Gale, 1981), decrease, or remain unaffected (Epron *et al*., 1999; Keiper *et al*., 1998; Koyro *et al*., 2006). Consequently, variations in respiration could be key in explaining small differences in yield among salt-stressed crop varieties. For instance, a susceptible wheat variety (*Triticum aestivum*) might show a marked increase in respiration rate under salt conditions, while a tolerant variety maintains respiratory homeostasis. This suggests that the tolerant variety directs less fixed carbon to respiration and more to growth (Kasai *et al*., 1998). Several studies have shown that salt-stressed plants exhibit changes in the ratio of CO_2_ produced to O_2_ consumed, implying that mitochondria may oxidise different substrates during salt stress (Acosta-Motos *et al*., 2017). The type of substrates respired significantly influences the plant’s overall metabolic balance, especially in regulating growth, compatible solute formation, and energy production.

There is a strong inverse relationship between the development rate and the amount of daily fixed carbon that must be respired (Hauben *et al*., 2009). Mitochondrial respiration is essential, and the benefit of a high respiration rate is that more ATP is generated, providing energy for processes such as ion exclusion, compatible solute synthesis, and detoxifying ROS (Munns and Tester, 2008). However, if mitochondrial respiration is hindered, ROS production increases and may surpass detoxification capacity (Jacoby *et al*., 2010). The ability of plants (such as wheat and rice) to exclude Na^+^ ions is closely linked to their root respiration rate (Malagoli *et al*., 2008). Evidence also suggests that maintaining respiratory homeostasis in the shoot is associated with salt tolerance (Jacoby *et al*., 2013). Nonetheless, confirming whether this is true across a broader range of species remains to be investigated. Species with higher energy-use efficiency may require less increased respiration for defence under salt stress, thereby reducing ROS production. Yet, photorespiration rates generally rise under salinity stress, adding to the ROS burden (Voss *et al*., 2013).

The TCA cycle links respiratory metabolism with carbon and nitrogen metabolism by producing substrates for mitochondrial respiration, which are central to the plant metabolic network (Nunes-Nesi *et al*., 2013). Several alternative pathways or shunts of TCA cycle metabolism have been reported in plants, including the malate-pyruvate pathway catalysed by malic acid, β-oxidation, and the GABA shunt. The gamma-aminobutyric acid (GABA) shunt was first identified in potatoes (*Solanum tuberosum*) more than 70 years ago (Dent *et al*., 1947); however, its role in metabolism is still being refined. The GABA metabolic pathway (GABA shunt) bypasses two steps of the TCA cycle, oxidising 2-Oxoglutarate to succinate, by circumventing the activities of 2-Oxoglutarate dehydrogenase and Succinyl-CoA synthetase (Carillo, 2018; Che- Othman *et al*., 2017). GABA abundance is primarily regulated through its synthesis rate (Fait *et al*., 2005). However, increases in GABA are also observed in *Arabidopsis* GABA- T- deficient mutants under salinity stress, indicating that its concentration is also controlled by its degradation rate (Renault *et al*., 2010). They reported that the GABA-T mutants exhibited an ultra-sensitive reaction to salinity and an increase in GABA levels in response to salt stress, which they suggested could result from GABA-T acting in reverse, catalysing the conversion from succinate semialdehyde (SSA) to GABA (Akçay *et al*., 2012). GABA and proline levels increase in response to salinity and other abiotic stresses in wheat (Al-Quraan *et al*., 2013; Che- Othman *et al*., 2019), maize (Wang *et al*., 2017), soybean (Xing *et al*., 2007), tobacco (Allan *et al*., 2008), sesame (Bor *et al*., 2009), and barley (Widodo *et al*., 2009). These two metabolites can be rapidly synthesised to provide cellular stress protection, primarily as osmolytes, ROS scavengers, or alternative metabolic shunts (Carillo *et al*., 2008). Simultaneously, several glutamate metabolites that serve as precursors for proline and GABA also increase under stress conditions (Carillo, 2018; Liu *et al*., 2011).

The synthesis of GABA from glutamate via glutamate decarboxylase leads to an energy reduction and releases CO_2_. This process helps the Calvin cycle stay active at lower membrane potential within the photosynthetic electron transport chain, thus reducing ROS formation and photodamage. In fact, inducing a functioning GABA shunt is crucial to limit ROS buildup (Renault *et al*., 2010). One study reports that GABA levels decrease in tobacco plants exposed to 500 mm NaCl, indicating salinity stress (Zhang *et al*., 2011), however, this may be due to salt shock from the sudden application of high NaCl concentrations. The GABA shunt activity depends on GAD activity, which is regulated by pH and calcium ion levels (Che-Othman *et al*., 2017). Enhanced GAD and GABA shunt activity increases respiration in salt-tolerant soybeans treated with CaCl_2_ (Yin *et al*., 2015). High GAD expression under salinity suggests GABA’s importance during wheat germination and development. This induction prompts a shift to an alternative metabolic pathway in the respiratory machinery, balancing carbon and nitrogen metabolism under abiotic stresses. Salt stress also increases levels of two other GABA shunt metabolites, glutamate and alanine (Che-Othman *et al*., 2019), as well as the activity of four GABA metabolic enzymes (Renault *et al*., 2010). Activation of the GABA shunt modulates glutamate levels, which can be significantly impacted by GAD overexpression, since GABA acts as a temporary nitrogen storage metabolite (Forde and Lea, 2007).

Metabolite and protein changes in the salt-treated plant respiratory system have been explored in several studies, and multiple correlative changes have been observed (Banaei-Asl *et al*., 2016; Che-Othman *et al*., 2019; Jacoby *et al*., 2011). However, only a few of these changes have been characterised at the mechanistic level, limiting the potential targets for breeding. Some metabolites, such as GABA, are proposed to have dual roles as alternative respiratory substrates and signalling molecules (Bandehagh and Taylor, 2020). The salinity-induced changes in metabolites involved in respiration may be crucial for plant salt tolerance, acting as sources of energy and carbon/nitrogen, or adjusting the osmotic potential within cellular compartments under salinity stress (Kosová *et al*., 2013). They can be synthesised *de novo* in response to salt stress or transported from other subcellular locations, but studying these remains challenging because of the mitochondrial context within the whole-cell metabolome. Organelle-targeted proteomics offers deeper insights into compounds that interact with, stay in, or are altered by stresses. Analysing enriched mitochondrial fractions from various wheat cultivars (Jacoby *et al*., 2016; Jacoby *et al*., 2013) and other crops, along with stress treatments, has enhanced our understanding of metabolic responses to salt exposure. In this study, we examined how salt stress affects barley respiratory metabolism, a crop with moderate salt tolerance compared to our previous research on wheat, revealing both shared and divergent responses to salt exposure. We found that barley mainly depends on the classical TCA cycle under salt stress, indicated by increased pyruvate oxidation and downstream metabolites. Meanwhile, the GABA shunt seems to play a limited role, unlike in wheat, where GABA shunt activity is heightened. The decreased levels of 2-oxoglutarate, succinate, arginine, and ornithine in barley support the idea that the GABA shunt plays a minor part in its salinity tolerance.

## Materials and Methods

### Plant growth

Barley plants (*Hordeum vulgare* L. var. La Trobe) were cultivated in a controlled hydroponic system. Seeds were germinated in Petri dishes with filter paper soaked with Thiram (dimethylcarbamothioylsulfanyl N, N-dimethylcarbamodithioate) solution (1.4 g/L). After 7 days, the seedlings were moved to a flood-drain hydroponic system containing a half-strength modified Hoagland solution ( 1 mM NH_4_NO_3_; 0.5 mM KH_2_PO_4_; 0.25 mM CaCl_2_; 50 μM KCl_2_; 0.5 mM MgSO_4_; 25 μM H_3_BO_3_; 2 μM MnSO_4_; 2 μM ZnSO_4_; 0.5 μM CuSO_4_; 0.5 μM Na_2_MoO_4_ and 0.5 mM Fe-EDTA) and pumps were turned off for 15 minutes every 30 minutes, for aeration. Plants were cultivated in a laboratory growth chamber with a 16/8 hour light/dark cycle, a light intensity of 500 µmol m^−2^ s^−1^, a day/night temperature of 28/22 °C, and a constant humidity of 65 per cent. The Hoagland solution was changed every 7 days. Salt treatments began on the eighth day after emergence. Over four days, 25 mM increments of NaCl were added to the nutritional solution in the morning and evening to attain a final NaCl concentration of 200 mM.

### Measurements of photosynthesis and respiration

Carbon dioxide fluxes of the third leaf middle section were determined using a portable infrared gas analyser (LI-COR 6400XT; LI-COR Inc., Lincoln, NE, USA) with a 60 x 60 mm leaf chamber on day 11 after salt treatment. The measurement was started a few hours after the growth cabinet’s illumination period began. The LED light source in the leaf chamber was adjusted to photosynthetically active radiation (PAR) of 500 µmol m^−2^ s^−1^, and the flow rate through the leaf chamber was adjusted to 300 mol s^−1^. Respiration measurements were taken with a Q2 oxygen sensor (Q2 platform-Astec Global, Maarssen, The Netherlands) in sealed 850 mL tubes holding three 1 cm^2^ third leaf discs. At 3-minute intervals, the oxygen concentration was measured. The oxygen consumption slope was determined between 0.5 and 3 hours following the run’s commencement. To calibrate to 100% and 0% atmospheric oxygen, standard samples containing normal air and 100% N_2_ were employed. The oxygen partial pressure was determined to be 20.95 per cent of atmospheric pressure, and molar oxygen consumption rates were calculated using the ideal gas law (Scafaro *et al*., 2017).

### Biomass and elemental sodium content

Roots, shoots, and the third emerging leaf were separated and harvested on days 1, 4, 8, 11, and 15 after 200 mM NaCl addition. Fresh weight was recorded, and then they were oven-dried at 60°C.

For elemental analysis, oven-dried samples were ground into a fine powder, digested in 10 mL of 0.5 M HCl, and shaken for 2 days using a rotating shaker. The extracts were then filtered through a 0.45 µM syringe-driven filter into 50 mL glass vials. The sodium and potassium content of the filtrates was measured using a flame photometer (M410, Sharwood Scientific, Cambridge, UK).

### Mitochondrial isolation, protein extraction and digestion

Mitochondria were isolated from whole shoots according to the method of Jacoby *et al*. (2013) Kerbler and Taylor (2017) using differential centrifugation, followed by a PVP (0 to 4.4% (w/v)) and Percoll self-formed gradients. Mitochondrial proteins were precipitated and extracted following the method of Huang *et al*. (2014).

### MRM optimisation and targeted protein quantification

Peptide transitions were optimised following the approach described in Taylor *et al*. (2014) using trypsin-digested isolated mitochondrial extracts run on an Agilent 6430 triple quadrupole (QqQ) mass spectrometer with an HPLC Chip Cube source and a UHC (II) chip (Agilent Technologies, Santa Clara, CA, USA).

### MRM data analysis

Skyline (MacLean *et al*., 2010) was used to analyse raw MS data, and peptides with fewer than 3 reliable transitions were removed. MetaboAnalyst (https://www.metaboanalyst.ca) version 6.0 (Pang *et al*., 2024) was used to examine differences in protein abundance between mitochondria isolated from control and salt-treated plants. To eliminate biases between MS runs, the transitions’ peak intensities were log2-transformed and standardised across runs using the equalised median algorithm.

### Gas Chromatography-Mass Spectrometry (GC-MS)

Metabolites were extracted using the method developed by Shingaki-Wells *et al*. (2011). The samples were analysed using a GC 6890n gas chromatograph (fitted with a 7683B Automatic Liquid Sampler and 5975B Inert MSD quadrupole MS detector (Agilent Technologies, Santa Clara, CA, USA) based on the method of Howell *et al*. (2009). The abundance of each metabolite in each sample was evaluated relative to internal standards (D-Sorbitol-^13^C_6_ and L-Valine-^13^C_5_).

### Statistical Analysis

Metabolomic data were normalised to internal standards and analysed using the online tool MetaboAnalyst (https://www.metaboanalyst.ca) version 6.0 (Chong *et al*., 2019). Significant metabolites were determined using a t-test and False Discovery Rate correction in MetaboAnalyst. Heat maps, principal component analysis (PCA), and partial least squares discriminant analysis (PLS-DA) were performed using MetaboAnalyst. A random forest analysis was conducted to assess the significance of metabolites in stressed plants. Metabolites were also evaluated for their reliability as biomarkers in stressed plants using Receiver Operating Characteristic (ROC) analysis. The actual positive rate (sensitivity) and valid negative rate (specificity) were estimated at a 0.25 threshold, and all metabolites were ranked based on the area under the curve (AUC) values at a 95% confidence level. The metabolic pathway was surveyed using pathway analysis in MetaboAnalyst, with rice metabolic pathway databases serving as a reference for the global test algorithm, based on the KEGG metabolic database and pathway construction.

## Results

### Growth, photosynthesis, respiration, chlorophyll concentration, sodium and potassium content

The sodium and potassium abundance in the third leaf of control and salt-treated plants was analysed over 11 days to assess the concentration of both ions in barley leaves (Figure 1a and b). From day 1 to day 15, the leaf sodium content of salt-treated plants increased to around 3.6-fold greater than the control (Figure 1a), and the leaf potassium content of salt-treated plants was reduced by approximately half of the control (Figure 1b). The mean K^+^/Na^+^ ratio of salt-treated plants was approximately 13% of that of control plants from day 1 to day 15 (Figure 1c). The impact of the adsorption of both ions on biomass accumulation, chlorophyll concentration, photosynthesis, and respiration rate in the roots and shoots was investigated (Figure 2a-g). Root, shoot, and third leaf dry weight, chlorophyll concentration, and photosynthetic rate were considerably reduced by salt treatment (Figures 2a, b, c, e, and f). Interestingly, SPAD values (chlorophyll concentration) were initially higher in salt-treated leaves than in controls, but declined over time to levels lower than those of the controls. In contrast, the respiration rate (O_2_ consumption) was significantly greater than that of control plants (Figure 2g).

**Figure 1.**
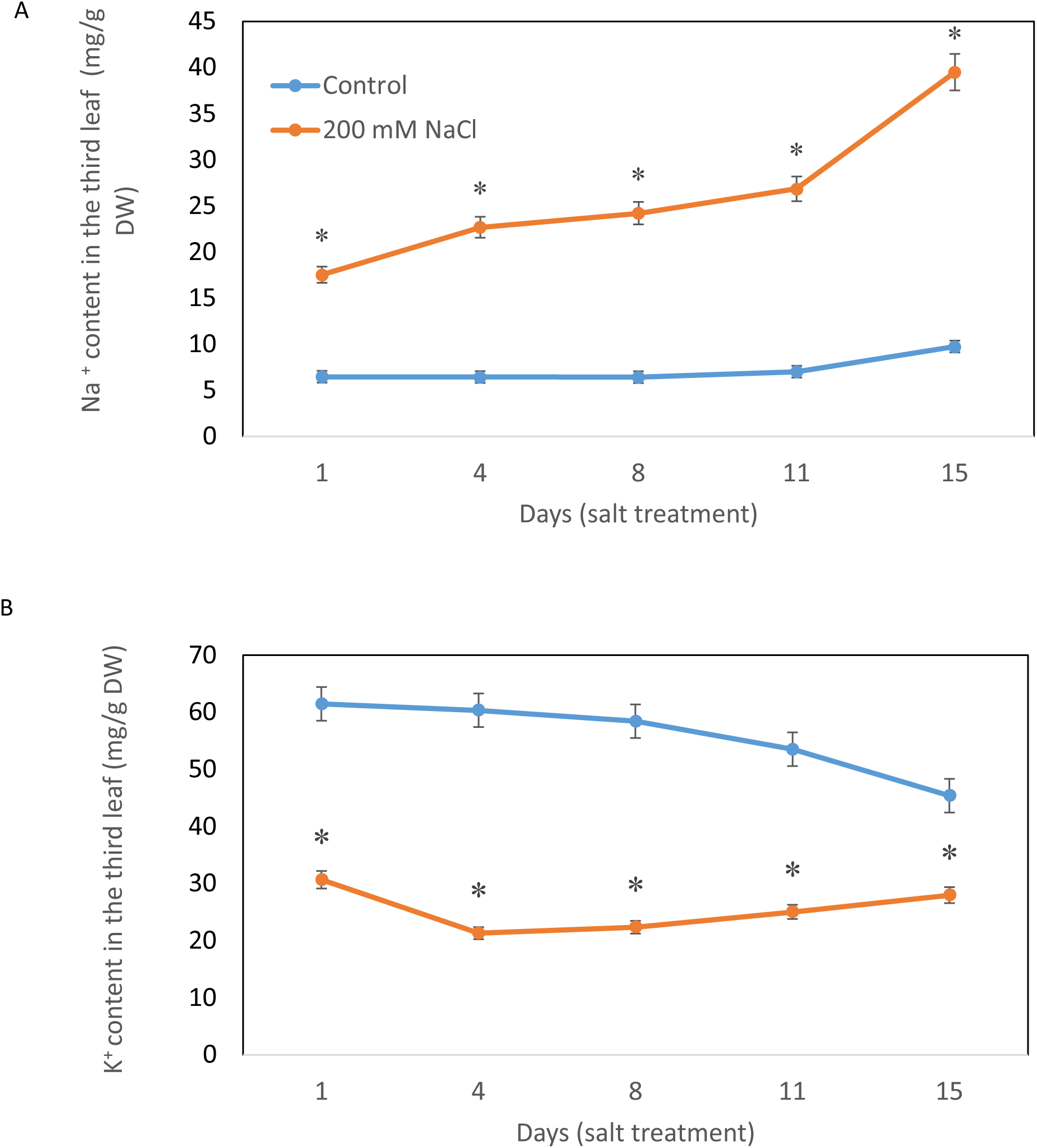
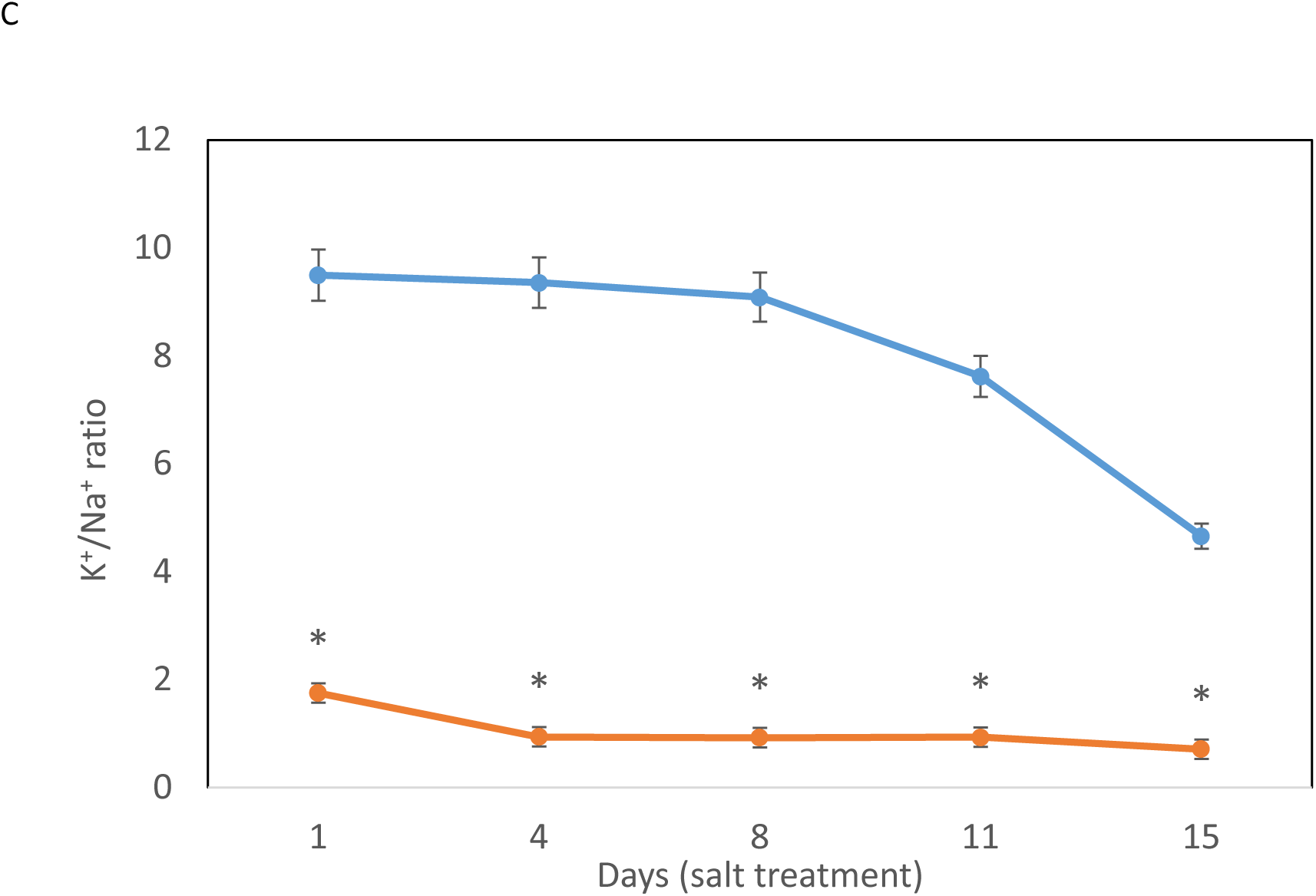
Ionic responses to salt stress of barley (*Hordeum vulgare* L. var La Trobe). (**a**) Sodium content, (**b**) Potassium content and (**c**) K^+^/Na^+^ ratio of barley leaf 3 at day(s) 1, 4, 8, 11 and 15 after 200 mM NaCl addition. The blue and red lines represent the control and salt-treated conditions, respectively. Error bars represent SEM. Stars indicate a significant difference between control and salt-treated plants (P<0.05), n=4.

**Figure 2.**
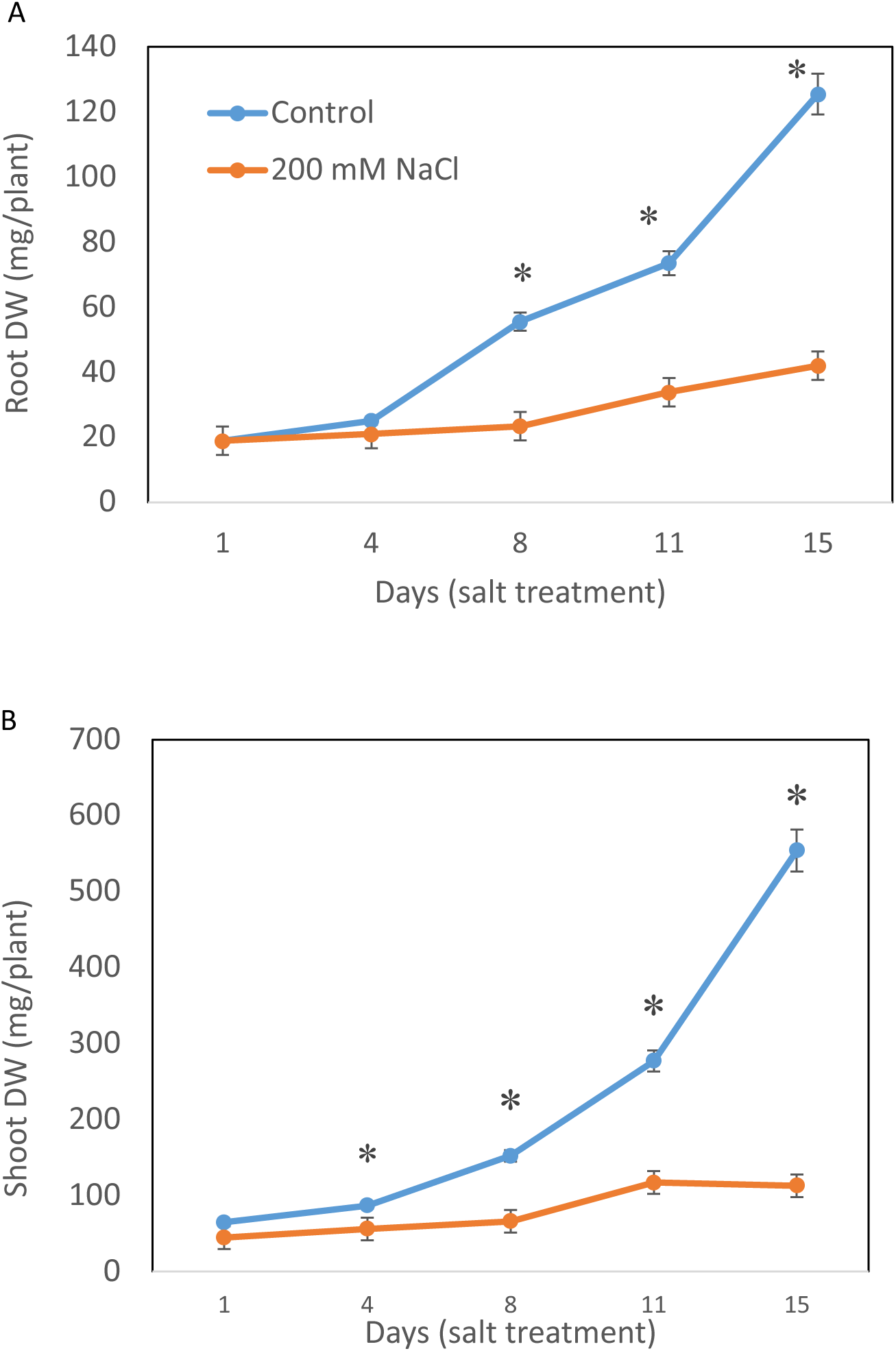
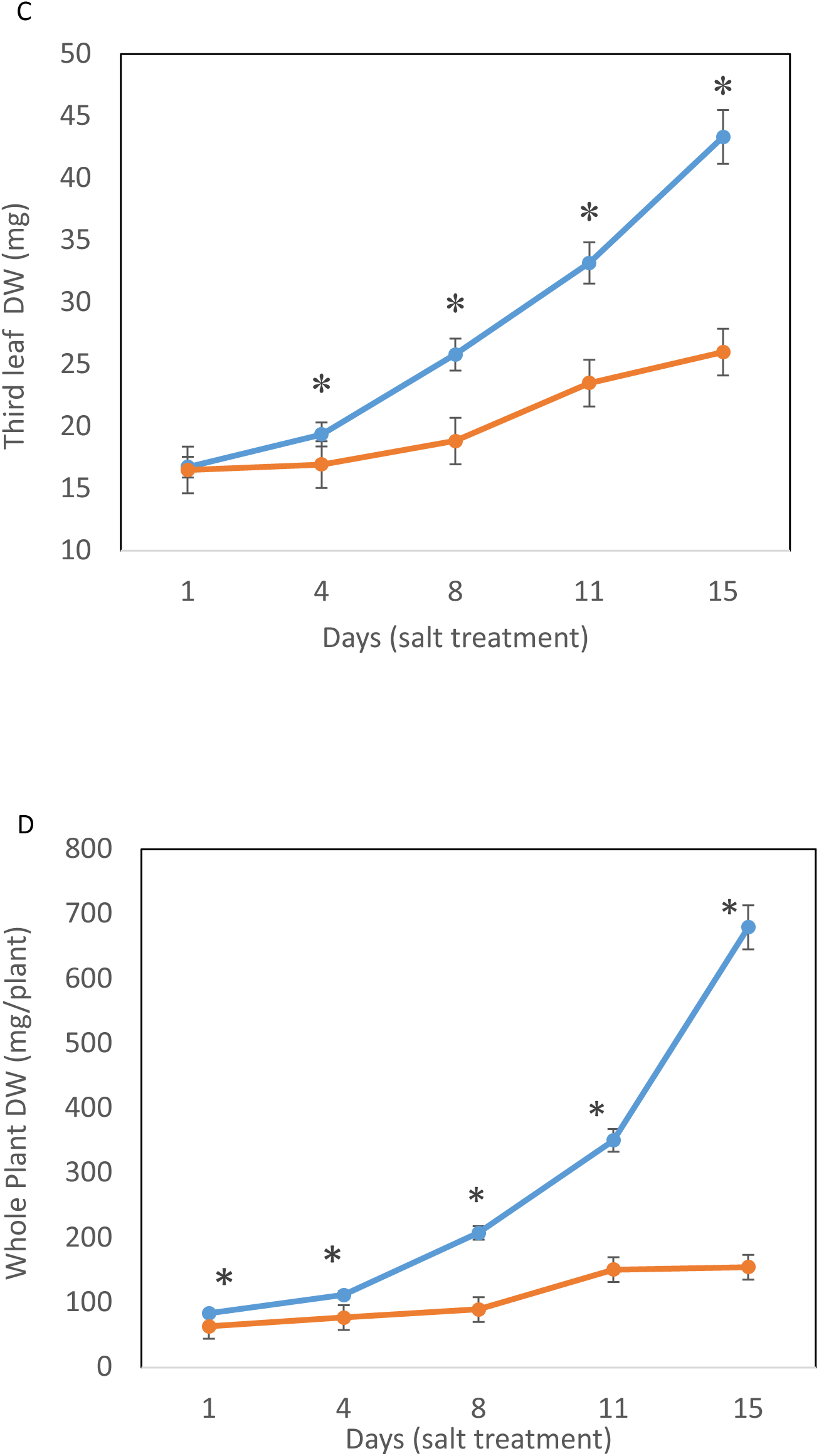
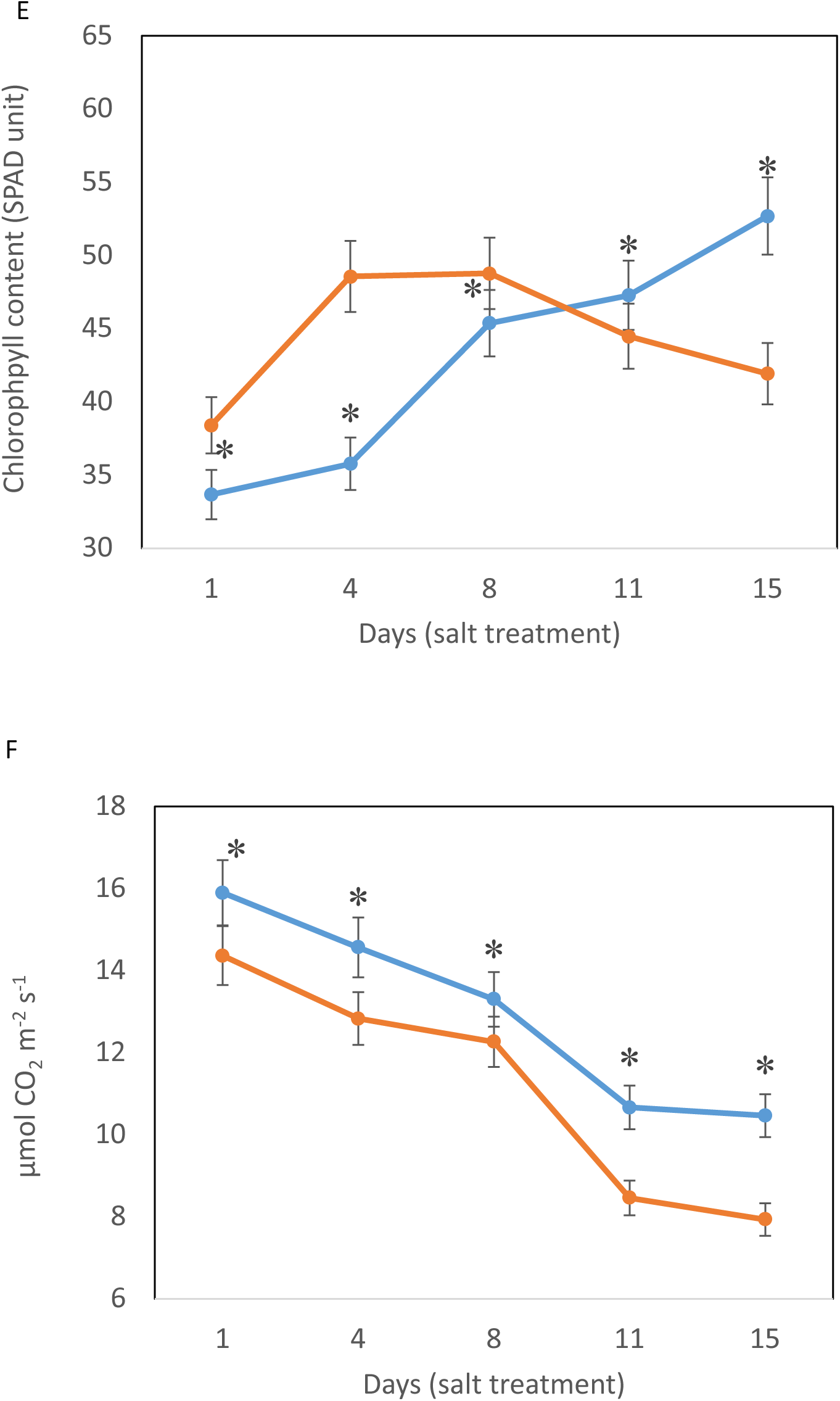
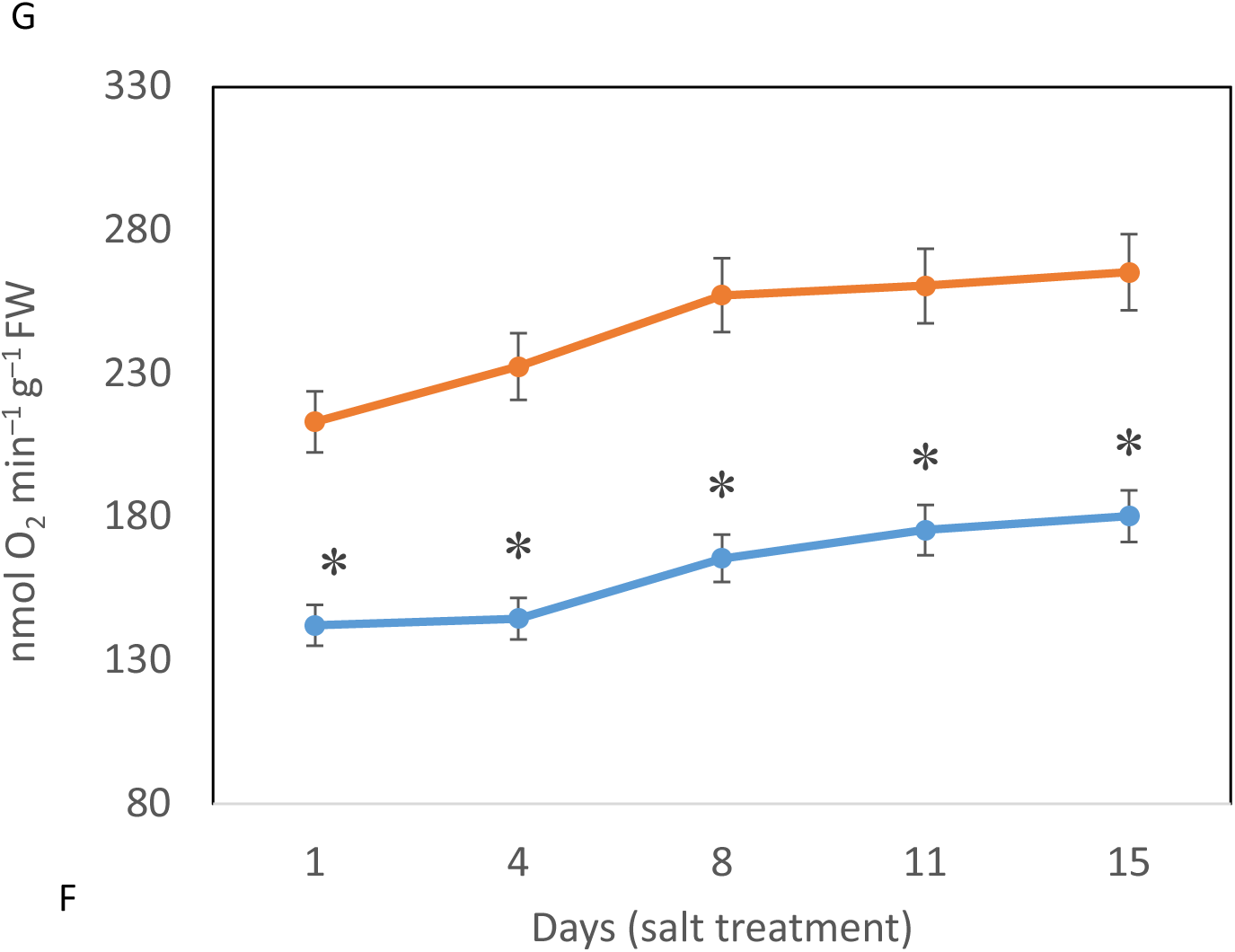
Physiological responses of barley to salinity stress. **(a)** Root dry weight (DW), **(b)** Shoot DW, **(c)** Third leaf DW, **(d)** Average biomass of barley plant, **(e)** Chlorophyll content (SPAD units), **(f)** Photosynthesis rate (µmol CO2 m^−2^ s^−1^), and **(g)** Respiration rate (nmol O_2_ m^−2^ g^−1^ FW) of barley in control and salt-treated plants (day 1, 4, 8, 11 and 15 after 200 mM NaCl addition). Error bars represent SEM. Stars indicate a significant difference between control and salt-treated plants (P<0.05), n=4.

### Metabolite changes following salt exposure

On day 11 of the salt treatment, the third leaf was collected from both control and salt-stressed plants, and metabolite profiles were analysed using GC-MS to investigate changes in metabolite abundance in barley leaves under salinity stress (Figure 3a). These changes indicate how salt inhibits enzymes in central carbon metabolism, suggesting a potential impact on overall plant performance. These metabolites were categorised into 13 major metabolic pathways. They included metabolites involved in amino acid metabolism (alanine, aspartate, glutamate, arginine), the TCA cycle, aminoacyl-tRNA biosynthesis, glyoxylate and dicarboxylate metabolism and arginine and proline metabolism (Figure 3b). The samples were grouped into two groups (control and salinity stress) using unbiased PLS-DA, which is presented as a score plot (Figure 3c). This shows a clear separation between control and salinity-treated samples, primarily along Component 1, which accounts for 97.7% of the variance. This separation indicates that salinity stress induces significant changes in metabolites, with both groups showing tight clustering, indicating a consistent and measurable impact on barley. Furthermore, The PLS-DA analysis identified several metabolites with variable importance in projection (VIP) scores greater than 1, indicating their significant contribution to the discrimination between control and salinity treatments. Among these, asparagine, proline, glutamine, and pyroglutamic acid displayed the highest VIP scores, with proline and asparagine particularly enriched under salinity stress. Glucose also showed a strong contribution, whereas other amino acids such as alanine, serine, glutamic acid, and tyrosine had lower but still notable impacts. (Figure 3d). Biomarker analysis revealed that most metabolites were highly reliable biomarkers, with Area Under the Curve (AUC) values above 0.9 (Supplementary Table 1).

**Figure 3.**
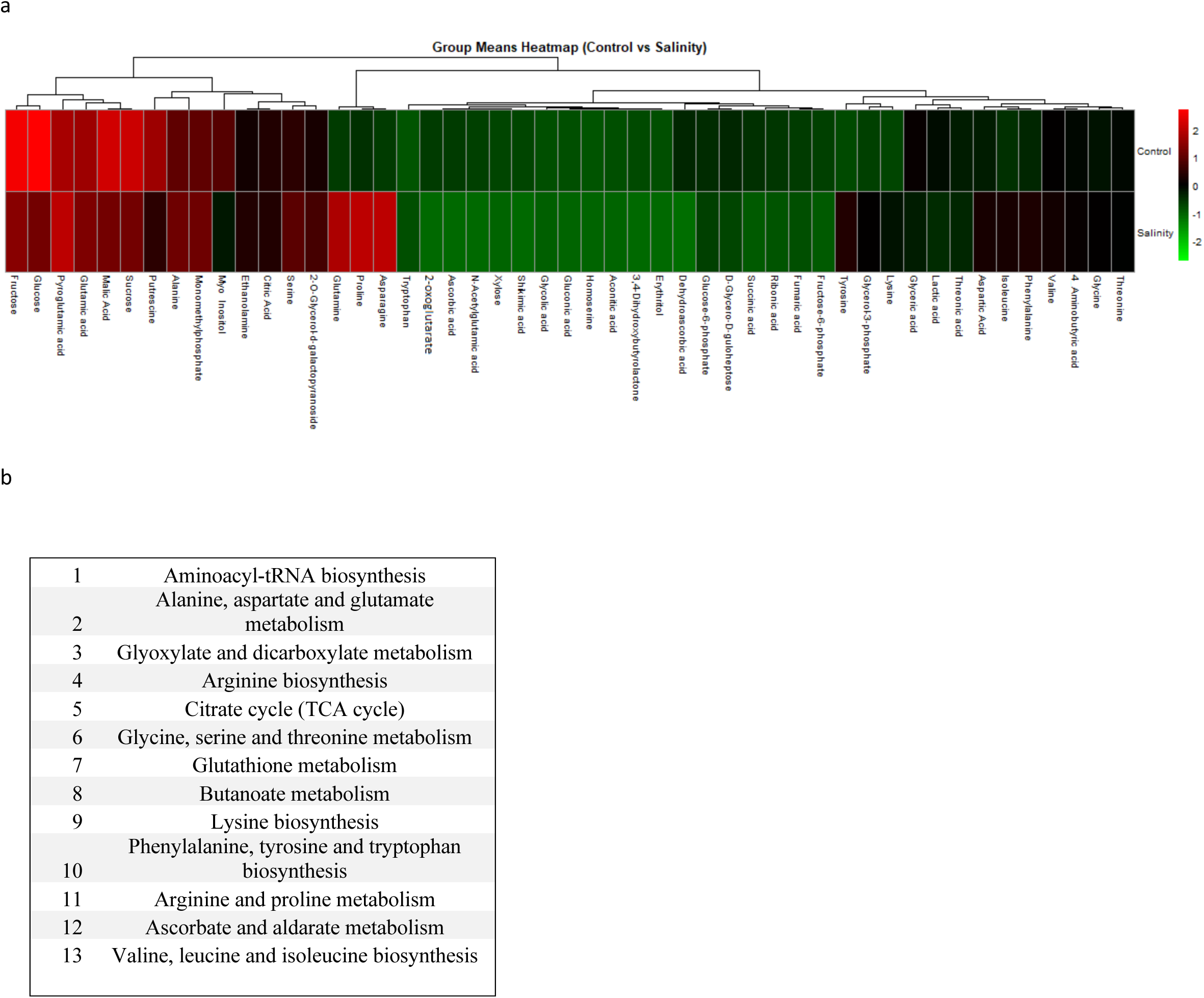
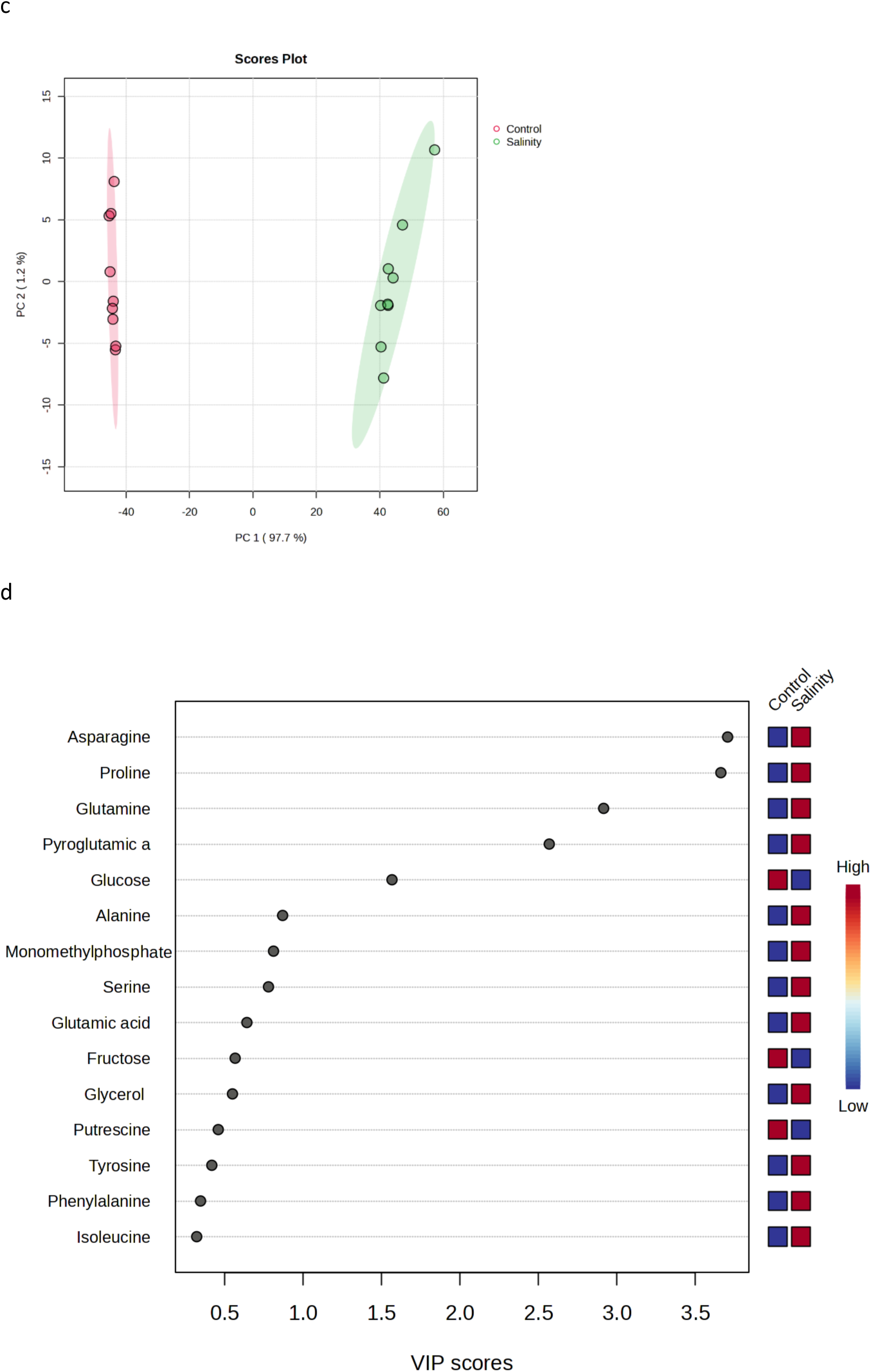
Targeted metabolomics analysis of barley leaf 3 on day 11 after 200 mM NaCl addition **(a)** Heat map of significant differential metabolites between control and salt-stressed plants. **(b)** Metabolic pathways to which differential metabolites are components at an FDR of 0.25. **(c)** Partial Least Squares – Discriminant Analysis of control (red) and salt-stressed plants (green) shown as a score plot. **(d)** Variable importance plot (VIP scores) from the PLSDA. a-f were generated using MetaboAnalyst 6.0.

The abundance of many amino acids increased significantly after salt stress; for example, asparagine accumulated more than 150-fold, while glutamine and proline accumulated 120- and 104-fold, respectively (Figure 4). The abundance of tyrosine, tryptophan, lysine, and glycerol all increased. Glucose-6-P, fructose-6-P, glycerol, guloheptose, and 2-O-glycerol-α-d-galactopyranoside all significantly increased, while other sugars detected in this study exhibited significant decreases. The measurement of organic acid abundance detected a notable reduction in dehydroascorbic acid (the oxidised form of ascorbic acid) abundance (> 78-fold) in salt-stressed plants compared to control plants. At the same time, N-acetyl glutamic acid, ascorbic acid, 2-oxoglutarate, shikimic acid, and succinate all exhibited substantial reductions in abundance. On the other hand, several organic acids showed a significant increase in abundance, including glycolic acid, aspartic acid, aconitate, pyroglutamic acid, citrate, and gluconic acid in the salt-treated plants (Figure 5).

**Figure 4.**
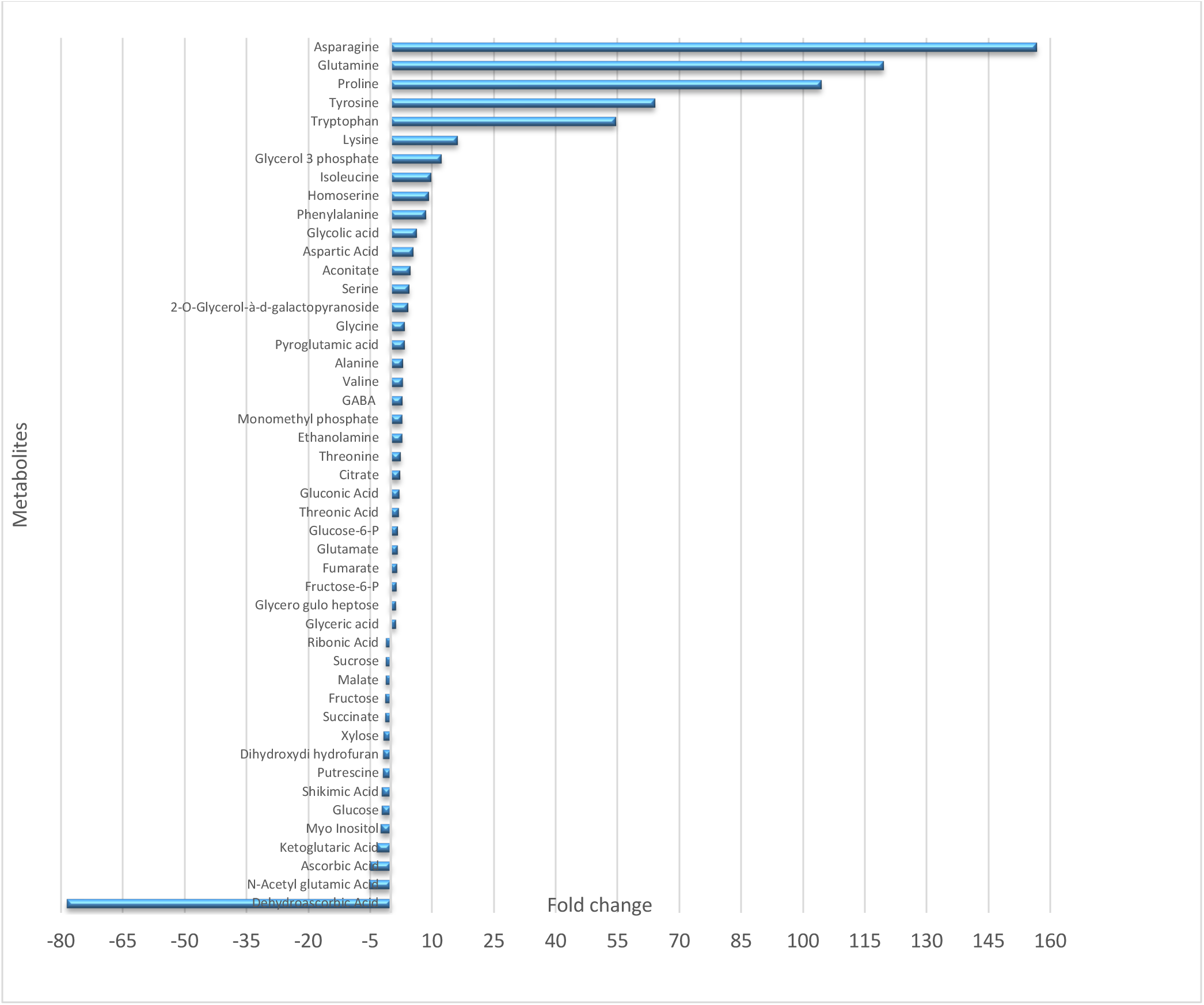
Changes in the abundance of metabolites of barley leaf 3 at day 11 after 200 mM NaCl addition. All metabolites show significant differences compared to the control, with P<0.05; n=4.

**Figure 5.**
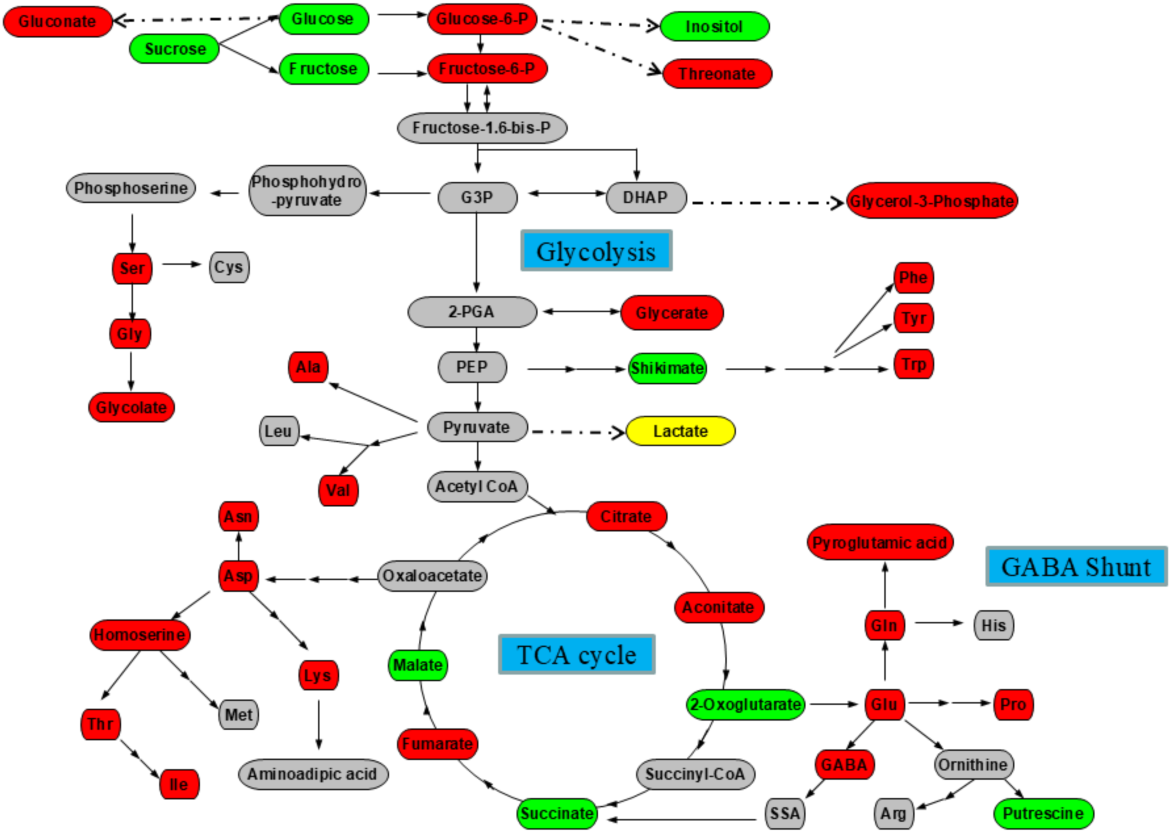
Summary of metabolic changes in metabolites of barley leaf 3 at day 11 after 200 mM NaCl addition relative to control conditions. Red boxes indicate up-regulation, and green boxes indicate down-regulation under salt stress. A yellow and grey box indicates unchanged and undetected metabolites, respectively. Abbreviations: G3P, glyceraldehyde 3-phosphate; DHAP, dihydroxyacetone phosphate; 1,3-BPG, 1,3-bisphosphoglyceric acid; PEP, phosphoenol pyruvate; PGA, phosphoglycerate. Amino acids are abbreviated to standard three-letter codes.

### Changes in the abundance of targeted proteins in response to salt exposure

Proteins were extracted from leaf three at day 11 of NaCl treatment from control and salt-stressed plants, and SRM mass spectrometry was performed to quantify a targeted set of proteins involved in the TCA cycle, mitochondrial membrane transport, and the GABA shunt to investigate changes in mitochondrial protein abundance in barley leaves under saline conditions. In response to salinity stress, seventy-five proteins showed a statistically significant (P=<0.01, log2 (fold change) >□1, fold change >□2) differential abundance (Figure 6a and Supplementary Table 2). Hierarchical clustering analysis was performed on the 75 proteins identified in the treatments, clearly distinguishing two main groups between the control and stressed plants (Figure 6a). Protein changes were further grouped into control and salinity-stress conditions using unbiased PLS-DA, as shown in a score plot with two components (Figure 6b). Of the 40 mitochondrial proteins detected by proteomic analysis, 12 respiration-related proteins with significant abundance changes and involvement in mitochondrial respiratory metabolism were selected for detailed analysis and presentation in Figure 7 and 8. These proteins included key enzymes of the electron transport chain, oxidative phosphorylation complexes, and TCA cycle components, all critical for mitochondrial energy production under salt stress conditions. Notably, adenylate kinase exhibited the highest increase in abundance following salt exposure (Figure 7), suggesting a pronounced activation of nucleotide metabolism. This upregulation likely reflects the cell’s need to maintain energy balance and support ATP/ADP turnover, which is critical for coping with osmotic stress and sustaining essential metabolic processes under salinity conditions. The selection criteria focused on proteins with established roles in respiratory metabolism, ensuring that the analysis captured the most functionally relevant changes in mitochondrial energy metabolism pathways. The remaining proteins, including structural, transport, and other mitochondrial proteins not directly involved in the respiratory process, were excluded from this focused analysis to highlight the specific metabolic adaptations in barley mitochondria in response to salt stress. This analysis revealed that the abundance of all targeted TCA cycle proteins (phosphoenolpyruvate carboxylase (PEPC), 3-hydroxyacyl-CoA dehydrogenase, ATP citrate synthase, and the succinate dehydrogenase flavoprotein) showed statistically significant increases in response to salt treatment (Supplementary Table 2 and Figure 7). There were significant increases in the abundance of pyruvate kinase, pyrophosphate-fructose 6-phosphate 1-phosphotransferase beta subunit, ornithine aminotransferase and sucrose synthase, at the same time, there was a decrease in the abundance of adenylate kinase (Figure 7, 8).

**Figure 6.**
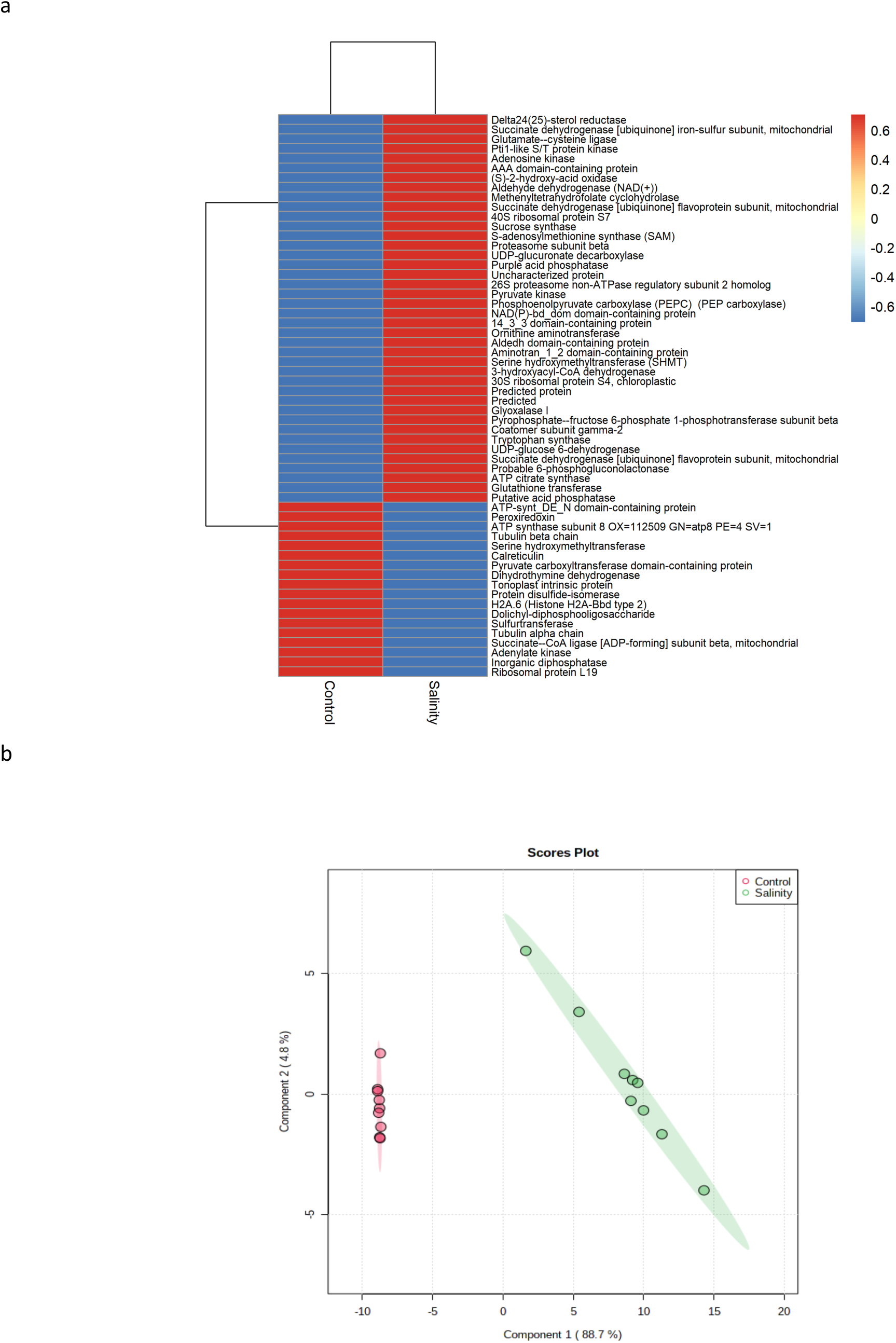
**(a)** Hierarchical cluster heat map of different detected proteins under salinity stress of barley leaves .**(b)** Unbiased PCA clustering of control (red colour) and salt-stressed plants (green colour) as represented by a score plot.

**Figure 7.**
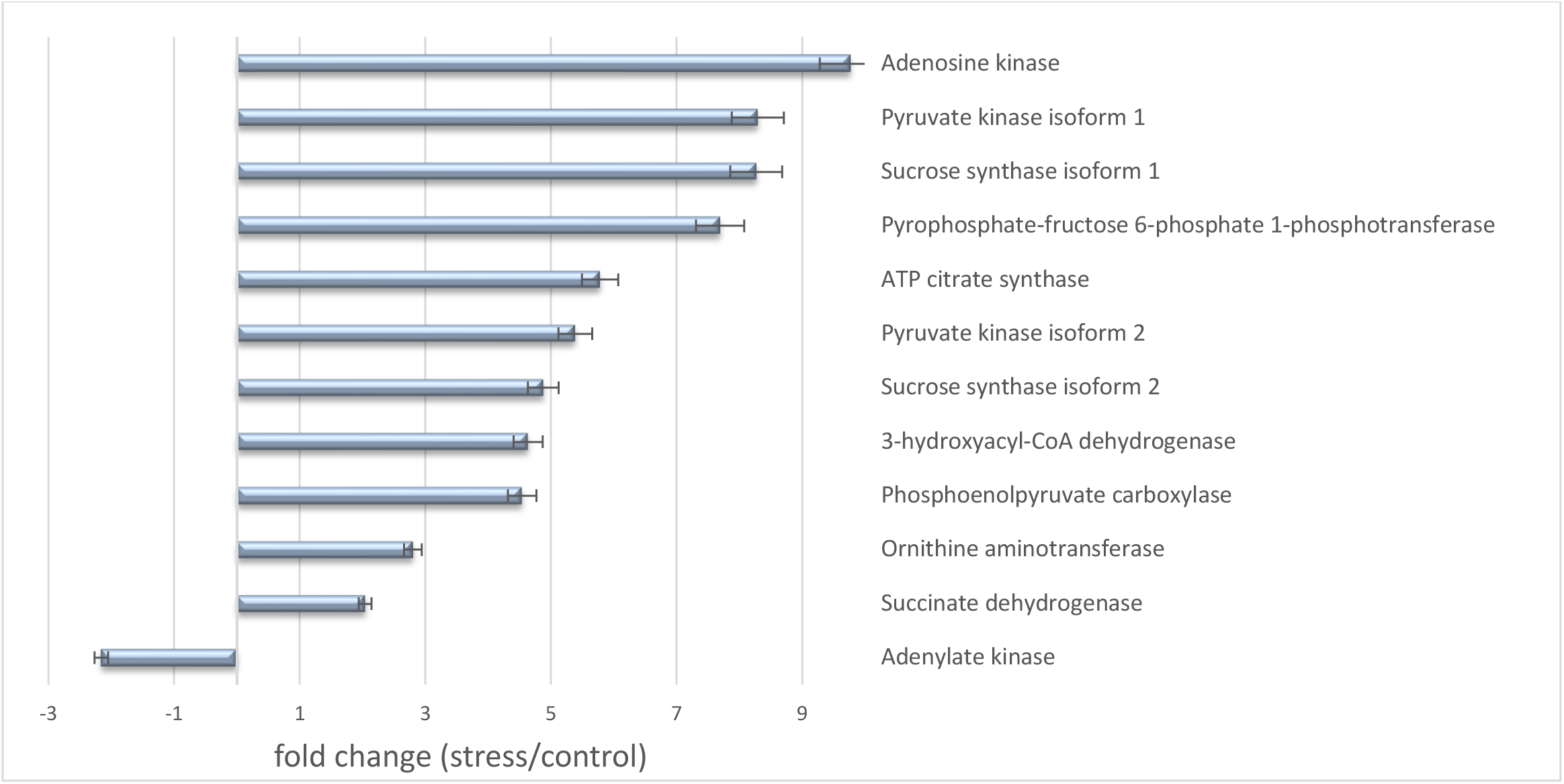
Significant changes in the abundance of targeted proteins in response to salt exposure. Comparisons were made between the control and 200 mM NaCl treatment in leaf 3 at day 11 after salt treatment. P=<0.01, log2(fold change) □>□1, fold change >□2, error bars represent SEM (n = 4).

**Figure 8.**
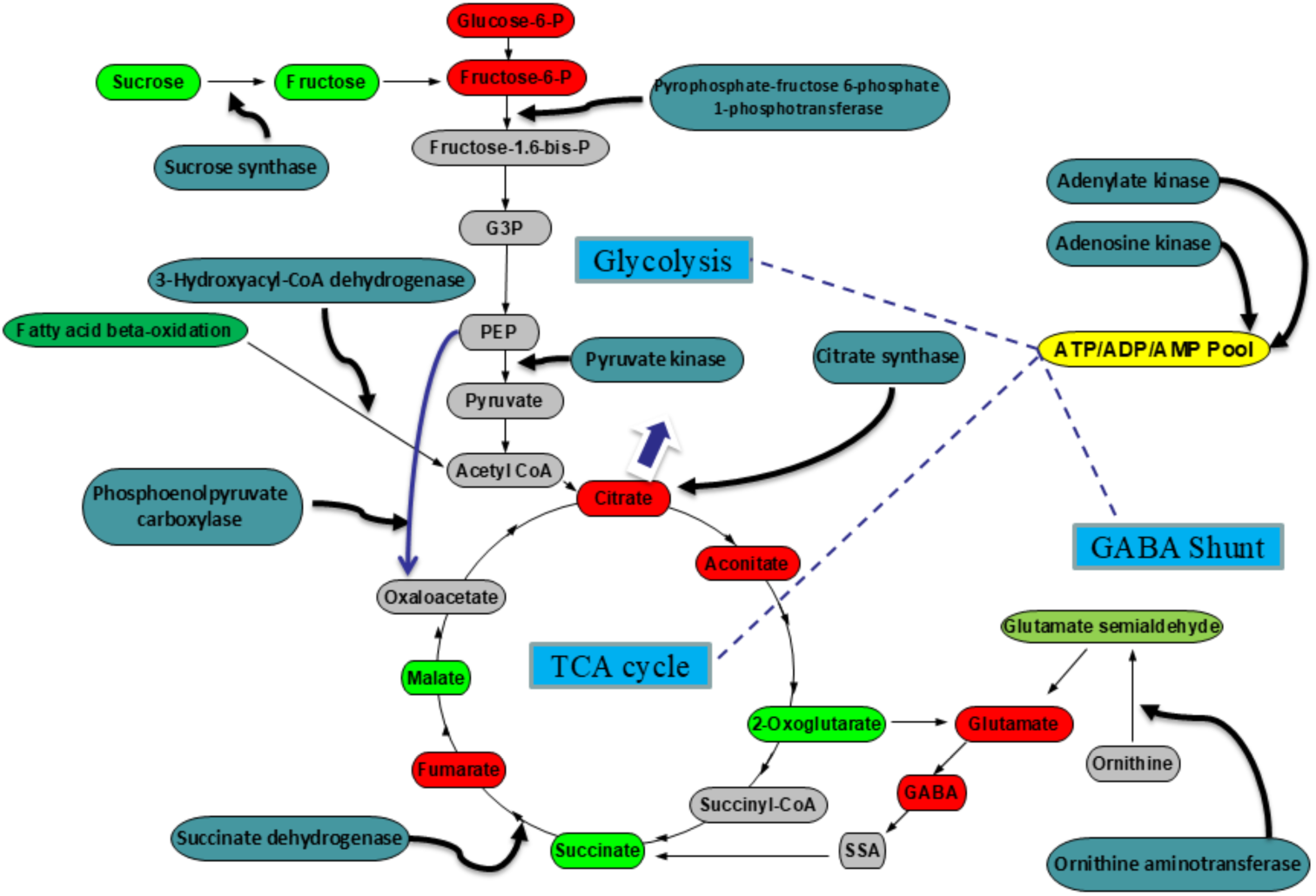
Summary of mitochondrial respiration–related proteins detected in barley (teal boxes), using the same samples employed for metabolite extraction.

## Discussion

When barley seedlings are exposed to salt, they accumulate significant amounts of sodium and show decreases in photosynthesis and biomass, indicating that salinity notably affects plant development and physiology. The combined effects of ions on their absorption are especially interesting under saline conditions. Ions like Na^+^ or Cl^−^ at high concentrations in the external solution are quickly taken up, leading to an excessive buildup in the tissue. Other ions, such as K^+^, may be hindered from entering the roots and being transported to the shoot via the xylem, thus lowering tissue concentrations (Cramer *et al*., 1991). Potassium levels and the K^+^/Na^+^ ratio decline as sodium content increases in stressed barley plants (Figure 1a and b). Although barley is regarded as a salt-tolerant crop, ionic and osmotic shocks can severely restrict its growth and development in salt-affected soils, with several responses linked to ionic stress. Under salt stress (200 mM NaCl), salt-tolerant barley genotypes produced more dry mass than salt-sensitive ones, and better tolerance was associated with a higher K^+^/Na^+^ ratio in the shoots (Mahlooji *et al*., 2018).

Measurements of respiration in the dark showed that O_2_ consumption increases in barley during salt stress and that salt is also transported from roots to leaves, causing ion-specific stress. This induces chlorosis and necrosis, along with decreased chlorophyll (Chl) content and reduced activity of key cellular metabolic pathways, including photosynthesis. These findings alone do not fully account for the changes in photosynthetic kinetics and respiratory rate observed in barley plants exposed to salinity. Che-Othman *et al*. (2019) showed that salt stress alters wheat mitochondrial respiration through an integrated analysis of metabolite levels, respiratory substrate utilisation assays, targeted quantification of protein abundance, and detailed enzymological measurements that explore the direct physicochemical effects of salt on enzymatic activity. Collectively, this revealed that salt treatment induces a coordinated set of metabolic changes in wheat cells, bypassing salt-sensitive steps in primary metabolism and reconfiguring respiratory metabolism, as evidenced by the metabolic fingerprint of salt-affected plants.

### Salinity stress alters the activity of the glycolysis pathway and pyruvate production for mitochondrial respiration

A possible mechanism for increased respiration involves heightened activity of components in respiratory energy metabolism, such as glycolysis, the TCA cycle, and the mitochondrial electron transport chain (ETC). An increase in O□ consumption in the dark signals a higher respiration rate and an adaptive response in energy metabolism. Glycolysis is a key metabolic pathway in plants that supports respiration by generating various intermediates, including ATP, NADH, and pyruvate. This study demonstrated that the enzyme pyruvate kinase, involved in glycolysis, was more abundant in salt-stressed barley leaves compared to control plants (Figure 7). Pyruvate kinase is a crucial regulatory enzyme that catalyses the irreversible substrate-level phosphorylation of ADP using phosphoenolpyruvate (PEP), producing pyruvate and ATP.

The change in abundance of pyrophosphate-fructose 6-phosphate 1-phosphotransferase beta subunit (PFP) indicated that this protein was more abundant in the leaves of salt-stressed plants compared to control plants (Figure 7). PFP catalyses the reversible interconversion of fructose-6-phosphate (Fru-6-P) and fructose-1,6-bisphosphate (Fru-1,6-P2), a rate-limiting step in controlling primary carbohydrate metabolism towards glycolysis. The variation in PFP abundance in salt-stressed plants suggests that this enzyme plays a key role in the adaptive response to salinity. Unlike ATP-dependent phosphofructokinase (PFK), PFP can utilise inorganic pyrophosphate (PPi) as an alternative phosphoryl donor, allowing plants to conserve ATP during stress. Since PPi is continually produced as a by-product of many biosynthetic reactions, its use by PFP offers a cost-effective mechanism to sustain glycolytic flux when ATP levels fall under salinity stress. The increased abundance of PFP in salt-stressed leaves reflects an adaptive metabolic strategy to maintain energy balance, boost carbon flow through glycolysis, and enhance stress resilience. This metabolic flexibility is especially vital in saline environments, where ATP availability is often limited. Consistent with this, studies in *Arabidopsis* have shown that PFP subunit expression is induced by salt and osmotic stress, and that PFP mutants exhibit significantly impaired growth under these conditions. Furthermore, altered PFP expression has been shown to affect plant performance in less-than-ideal environments, highlighting its role in stress adaptation (Duan *et al*., 2016; Lim *et al*., 2014; Lim *et al*., 2009).

### The TCA cycle responds to salinity stress

As a significant source of reductants to the electron transport chain (ETC), the tricarboxylic acid (TCA) cycle is a critical component of cellular ATP production. The proteins that form the TCA cycle have been identified, and their control has been intensively studied. However, the function and regulation of these proteins under salinity stress remain poorly understood. TCA cycle proteins, such as succinate dehydrogenase, are upregulated, indicating their significant role in providing vital energy for salinity tolerance. Succinate is oxidised by succinate dehydrogenase, and a lower abundance of this metabolite was observed under salinity stress, followed by an increase in succinate dehydrogenase abundance. In a flux control analysis of respiration of the TCA cycle, the antisense inhibition of succinate dehydrogenase reduced the respiration rate by approximately 15% despite a significant reduction in the activity of succinate dehydrogenase (60%) (Araujo *et al*., 2012) and a significant increase in the abundance of a particular isoform of succinate dehydrogenase subunit 2 has been observed following salt exposure in wheat plants (Che-Othman *et al*., 2019) and canola (Banaei-Asl *et al*., 2016). Paradoxically, another isoform of succinate dehydrogenase subunit 2 was shown to be less abundant after salt stress in wheat (Che-Othman *et al*., 2019), and the activity of this enzyme was reduced in mung bean roots, whereas it was stimulated in shoots under increasing concentrations of salt. (Saha *et al*., 2012)

Simultaneously, the activity of phosphoenolpyruvate carboxylase increased under salinity stress. Phosphoenolpyruvate carboxylase is a carboxy-lyase enzyme that catalyses the addition of bicarbonate to phosphoenolpyruvate, producing oxaloacetate and inorganic phosphate. Oxaloacetate results from the carboxylation of phosphoenolpyruvate (PEP), a four-carbon dicarboxylic acid source for the tricarboxylic cycle, with the phosphoenolpyruvate carboxylase gene family showing different expression patterns in Arabidopsis organs and in response to environmental stress. It was also observed that phosphoenolpyruvate carboxylase helps the plant adapt to salt and drought conditions. During salt stress, PEP carboxylase activity rose sharply in crops like maize developing leaves (Hütsch et al., 2016) and durum wheat (Fercha et al., 2013). The increase in PEP-carboxylase activity under salinity stress seems to compensate for reduced sink activity and may serve an anaplerotic function, supporting metabolite demands in the developing shoot tissues of young plants.

In our study, the expression of 3-hydroxyacyl-CoA dehydrogenase, an enzyme involved in β-oxidation that can produce acetyl-CoA, was significantly up-regulated by salinity stress, leading to increased acetyl-CoA entering the TCA cycle for energy production. Mitochondrial fatty acid β-oxidation is the process by which fatty acid molecules are broken down (Xu *et al*., 2017) in the cytosol of prokaryotes and the mitochondria of eukaryotes to produce acetyl-CoA, which then enters the TCA cycle as NADH and FADH_2,_ coenzymes used in the electron transport chain to generate ATP. The TCA cycle consumes acetate (in the form of acetyl-CoA), converting NAD^+^ to NADH, and is crucial for synthesising respiratory ATP by providing a reductant for the ETC. The β-oxidation pathway also occurs in peroxisomes and, similar to the GABA shunt, offers a way for cells to maintain carbon flow through primary metabolism when overall respiration rates are insufficient to meet energy demands. A metabolite profiling study in salt-treated soybean seedlings highlighted a significant role for the β-oxidation pathway (Zhang et al., 2016). Acetyl-CoA is converted into citrate, which is utilised in mitochondrial respiration. This increase in citrate production via β-oxidation has been observed in sweet potatoes under salinity stress (McLachlan *et al*., 2016). They noted a notable accumulation of lipids in the vegetative tissues of sweet potatoes under NaCl stress , alongside disrupted K^+^/na^+^ homeostasis, and exogenous stimulation of this pathway enhanced plasma membrane H^+^–ATPase activity and K^+^/Na^+^ balance(Yu et al., 2018). In this plant, isocitrate lyase and citrate synthase use the resulting acetyl-CoA to catalyse the entry of succinate and citrate into the TCA cycle.

Regulation of energy metabolism under salt stress is a crucial mechanism for salt tolerance, and ATP-citrate synthase is one of the most essential enzymes in respiration. The enzyme ATP-citrate synthase, also known as ATP-citrate lyase, catalyses the synthesis of acetyl-CoA and oxaloacetate from citrate and CoA (Coenzyme A). Because acetyl-CoA is a precursor in the biosynthesis of many chemicals in plants, distinct mechanisms for acetyl-CoA metabolism must exist in mitochondria (TCA cycle), plastids (*de novo* fatty acid synthesis), peroxisomes (β-oxidation), and the cytoplasm (biosynthesis of isoprenoids, flavonoids) (Pütter *et al*., 2017). Analysis of 2D-DIGE gels revealed that ATP-citrate synthase was differentially lower under salinity stress (Vítámvás *et al*., 2015). One unigene coding for ATP-citrate synthase was identified as up-regulated in wheat plants under salt stress. In our data, ATP-citrate synthase showed a 5.78-fold increase, suggesting that citrate breakdown could decrease the abundance of this metabolite. Interestingly, citrate increased in abundance following salt exposure, which must be produced for another part of metabolism. Several salinity studies show a lower abundance of several organic acids, most notably citrate, in wheat leaves (Che-Othman *et al*., 2019), barley leaves (Wu *et al*., 2013), and rice roots (Sanchez *et al*., 2008), indicating that these metabolites, which are immediately downstream of pyruvate in the TCA cycle, are inhibited by reduced pyruvate oxidation. Furthermore, increased citrate and aconitate concentrations reflect higher activity of acetyl-CoA-dependent organic acid synthesis, implying that mitochondrial pyruvate oxidation is impaired in barley leaves under salinity.

### Maintenance of mitochondrial respiration

The primary roles of adenosine kinase (ADK) and adenylate kinase (AK) were identified in grapevine under salinity stress as part of electron transport and membrane-associated energy conservation. Adenylate equilibration in the mitochondrial intermembrane space maintains respiration and regulates cytosolic metabolism. Although adenosine kinase is not a stress-responsive enzyme, it is vital for sustaining transmethylation processes by acting as a broad metabolic regulator to lower cellular free adenosine levels. Sugar beet and spinach exhibited higher ADK activity than control plants, whereas neither canola nor tobacco did under salt stress (Weretilnyk *et al*., 2001). This enzyme, a candidate protein for plant breeding related to salt stress, showed decreased expression after salt stress only in salt-sensitive cultivars (Dezong *et al*., 2017). Adenosine kinase plays a role in ATP metabolism and is crucial for plant stress acclimation, as active stress responses require substantial energy.

Adenylate kinase supports mitochondrial respiration and, in both directions of AK activity, utilises one Mg-complexed and one free adenylate as substrates. It is active in the mitochondrial intermembrane space but not in the matrix, where low ADP levels during vigorous respiration inhibit the respiratory rate. The matrix exports the ADP generated by AK in exchange for ATP, providing a channel for ADP recycling during respiration (Igamberdiev and Kleczkowski, 2006). It was shown that AK activity is an inherent characteristic of a key family of membrane transporters, enabling optimal gating of energetically neutral or downhill substrate fluxes across membranes (Randak and Welsh, 2005). This protein is upregulated in wheat genotypes during salt exposure (Faghani *et al*., 2015; Fercha *et al*., 2013).

### Sucrose degradation and glycolysis

Sucrose synthase is crucial for breaking down sucrose and providing energy. The only known enzymatic pathways for sucrose cleavage in plants involve invertases and sucrose synthase reactions (i.e., two hexoses instead of fructose + UDPG) for various metabolic processes. This enzyme’s role in sucrose import may include a dual ability to direct carbon towards polysaccharide formation and to facilitate an adenylate-conserving respiration mechanism. It can easily support an ATP-saving respiration pathway (Koch, 2004). For plant growth and stress tolerance, respiratory metabolism is essential. Plants exhibit several unique features of respiratory metabolism, such as multiple entry points for sucrose and starch, pyrophosphate recycling, and ATP-dependent fructose-6-phosphate phosphorylation (Stitt, 1998). The glycolysis pathway converts glucose (derived from starch and sucrose) into pyruvate. The TCA cycle then uses pyruvate as a carbon source to generate reducing equivalents like NADH and FADH_2_ for use by the ETC. An increased flow of carbon from glycolysis results in higher production of NADH, FADH_2,_ and ATP via the TCA cycle. Oxidation of one sucrose molecule through glycolysis yields 4 ATP, 4 molecules of pyruvate, and 4 NADH (Munns *et al*., 2019).

In this study, a reduction in saccharide levels such as glucose, fructose, xylose, and sucrose has been observed in some plants exposed to salinity, including Arabidopsis, *Lotus japonicus*, rice (Sanchez *et al*., 2008), *Hordeum spontaneum, Hordeum vulgare* (Wu *et al*., 2013), and wheat (Che-Othman *et al*., 2019). Monosaccharides actively contribute to energy supply and osmotic regulation in plants. A significant decrease in sucrose levels was noted in this experiment (Figure 4), likely indicating sucrose consumption. Metabolomics studies have shown that glycolysis and sucrose metabolism depend on the timing and intensity of stress and salinity, representing long-term responses to salinity in Arabidopsis. During prolonged salt exposure, glycolysis and sucrose metabolism are co-induced, resulting in a concurrent reduction in the methylation cycle (Urano *et al*., 2010). It has been reported that there are two mechanisms for sucrose degradation: the first involves invertase activity, followed by classical glycolysis of hexose sugars, while the second begins with sucrose synthase (Huber and Akazawa, 1986).

### A key glutamate-metabolising enzyme is ornithine aminotransferase

The ornithine aminotransferase (OAT) enzyme facilitates the conversion of ornithine into pyrroline-5-carboxylate (P5C). P5C can then be transformed into glutamate and proline, which were significantly upregulated by salinity stress in this study. In salt-stressed barley plants, the ornithine pathway and the glutamate system likely contribute to proline synthesis through increased OAT activity. The rise in OAT protein (Figure 7) appears to drive the salt-induced increase in OAT activity. The proline metabolic intermediate P5C has been shown to directly suppress mitochondrial respiration in budding yeast (Nishimura *et al*., 2012), whereas in this experiment, mitochondrial respiration activity was stimulated. Besides its roles in proline biosynthesis and arginine catabolism, OAT also participates in stress-induced cytoplasmic proline accumulation, programmed cell death (PCD), and non-host disease resistance in plants (Anwar *et al*., 2018). Proline is involved in the production of 2-oxoglutarate, a key intermediate in the TCA cycle.

Glutamate serves as the source of various biosynthetic structures, including ammonia donors for the synthesis of glutamine and ornithine (Carillo *et al*., 2008). The ammonia group is transferred to oxaloacetate by glutamate via aspartate aminotransferase, producing aspartate, which is a precursor for synthesising asparagine, lysine, methionine, and isoleucine (Petrov *et al*., 2015). Additionally, the GABA shunt pathway begins with the decarboxylation of glutamate to form GABA and CO_2_ in the cytosol. In this study, asparagine, glutamine, and proline showed the most significant accumulation under salt stress (Figure 4). The abundance of asparagine, which can contribute to the synthesis of oxaloacetate, may originate from various sources, including protein degradation and *de novo* synthesis by assimilating N into C skeletons derived from glycolysis and the TCA cycle (Gilbert *et al*., 1998).

In addition to succinate, proline is another vital metabolite. Its accumulation and breakdown under environmental stress can provide reductants that support mitochondrial electron transport and generate ATP (Szabados and Savouré, 2010). The TCA cycle also plays a key role in supplying reductants to the mitochondrial electron transport chain. The levels of compatible solutes, such as proline (104.4-fold increase, Figure 4), glycine (3.4-fold increase, Figure 4), and putrescine (1.9-fold decrease, Figure 4), which sustain respiratory metabolism, rose in many plant species. Although GABA’s primary source is GAD activity (2.8-fold increase, Figure 4), it can also be derived from putrescine, which participates in glycine betaine biosynthesis (Shelp *et al*., 2012) and proline under oxidative stress (Signorelli *et al*., 2015).

### The GABA shunt pathway is less important in Barley salinity tolerance

Che-Othman *et al*. (2019) reported that an increased abundance of succinate and 2-oxoglutarate supports the induction of the GABA shunt under salinity, and increased GABA shunt activity offers a potential explanation for the heightened respiration rate observed in salt-treated wheat leaves despite the reduced potential for pyruvate oxidation by the TCA cycle. In this study, both succinate and 2-oxoglutarate showed decreases in abundance under salt stress (Figure 4). Therefore, it can be concluded that the GABA shunt is less crucial for barley’s salinity tolerance. Succinyl semialdehyde dehydrogenase (SSADH), glutamate dehydrogenase (GDH), and other enzymes related to the GABA shunt pathway were not detected under salt exposure in barley mitochondrial samples. GDH and SSADH are components of the GABA shunt pathway that can provide an alternative carbon source for the TCA cycle (Bouche and Fromm, 2004). The GABA shunt pathway yields succinate as a product. Hence, the increase in 2-oxoglutarate and succinate levels could serve as markers of GABA shunting, which bypasses the 2-oxoglutarate dehydrogenase step in the TCA cycle (Fait et al., 2008). The rise in these metabolites in wheat (Che-Othman et al., 2019) links respiration with nitrogen assimilation and carbon balance under salinity. However, GABA concentration showed different trends in wheat and barley crops. Elevated succinate and 2-oxoglutarate levels in wheat may relate to increased activity of SSADH and GDH in the GABA shunt. Additionally, arginine and ornithine were not detected in this study. The accumulation of both metabolites aligns with the induction of the GABA shunt. Therefore, it can be concluded that the GABA shunt is of lesser importance in barley’s salinity tolerance.

### Overview of metabolite changes in TCA cycle induction

Increased citrate and aconitate concentrations suggest an enhanced capacity for acetyl-CoA-dependent synthesis of these organic acids. In the TCA cycle, both metabolites are directly downstream of pyruvate. Therefore, we hypothesise that when barley leaves are exposed to salt stress, mitochondrial pyruvate oxidation increases due to a higher abundance of metabolites downstream of pyruvate and enhanced pyruvate oxidation. Our results contrast with those of Che-Othman *et al*. (2019). Analysis of organic acid levels in wheat plants showed a decrease in citrate and aconitate in salt-stressed plants and inhibited pyruvate oxidation. Additionally, pyruvate synthesis is connected to alanine, which is fed into the TCA cycle via the reverse reaction of alanine aminotransferase (Diab and Limami, 2016). However, a reduction in alanine has been observed in both wild and cultivated barley under salinity stress (Wu *et al*., 2013), and the increase in alanine in this experiment could be attributed to the induction of pyruvate metabolism.

A comprehensive assessment of rice root responses to salt stress revealed significant degradation of TCA cycle intermediates, including malic, 2-oxoglutaric, and shikimic acids. An increase in some amino acids followed this degradation (Zuther *et al*., 2007). Many organic acids related to the TCA cycle in flower and pod tissues, including isocitrate and aconitate, were also increased by salinity in the desi chickpea cultivar, cv Rupali (salt sensitive), but not in the salt-tolerant cv Genesis 836 (Dias *et al*., 2015). In other words, some salt-sensitive plant lines increasingly use these metabolites as precursors for *de novo* amino acid biosynthesis under salinity stress. In other reports, the metabolic responses of barley leaves (Widodo *et al*., 2009), barley roots (Wu *et al*., 2013), and soybean (Zhang *et al*., 2016) to salinity saw an increase in TCA cycle intermediates such as aconitate and citrate, suggesting the TCA cycle plays a key role in salinity tolerance. These findings align with our experiments. Conversely, lower levels of aconitate and citrate have been reported in wheat (Che-Othman et al., 2019), barley leaves (Wu *et al*., 2013), and rice roots (Sanchez *et al*., 2008). Vanlerberghe and McLntosh (1996) reported that citrate and isocitrate, two TCA cycle intermediates, influence mitochondrial retrograde signalling in response to modifications or perturbations within the organelle, transmitting signals (retrograde signalling) to the nucleus to trigger specific expression of nuclear-encoded genes. A significant decrease in dehydroascorbic acid (78-fold) was observed in this study (Figure 4). Ascorbic acid’s reversible oxidation to dehydroascorbic acid may function as a redox buffer during plant respiration. Bartoli *et al*. (2006) reported that the mitochondrial electron transport chain exerts coordinated control over redox pathways involving ascorbic acid capacity, and the plant’s ability to synthesise and accumulate ascorbic acid is modulated by mitochondrial electron transport capacity and components.

## Conclusion

While further experiments are needed to clarify how salinity stress affects respiratory control in plants, we demonstrate that salinity triggers mitochondrial respiration and impacts carbon and nitrogen metabolism. Glycolysis and sucrose metabolism are fueled by decreased levels of metabolites like glucose, fructose, and xylose, along with high levels of sucrose synthase and pyruvate kinase. This suggests that glycolysis becomes more active under stress, consuming phosphoenolpyruvate (PEP) to produce pyruvate and ATP. Metabolite analysis shows activation of the TCA cycle, with increased utilisation of pyruvate evidenced by higher levels of aconitate and citrate. The rise in these metabolites indicates enhanced pyruvate oxidation and increased capacity for pyruvate processing through the TCA cycle in barley. Succinate, which is oxidised by succinate dehydrogenase, initially showed lower levels, but subsequently, an increase in succinate dehydrogenase abundance and respiration rate was observed in this experiment. Phosphoenolpyruvate carboxylase activity produces oxaloacetate from phosphoenolpyruvate, supplying carbon to the TCA cycle. Additionally, activity of 3-hydroxyacyl-CoA dehydrogenase may facilitate greater entry of acetyl-CoA into the TCA cycle. We hypothesise that under salt stress, mitochondrial pyruvate oxidation is induced in barley. Elevated levels of metabolites downstream of pyruvate suggest that the TCA cycle is crucial for salinity tolerance. Our findings also indicate that the GABA shunt is not vital in barley, as the decrease in 2-oxoglutarate and succinate levels could mark reduced GABA shunting activity. Furthermore, arginine and ornithine were not detected in this study, although their accumulation would be consistent with GABA shunt induction. Consequently, we may conclude that GABA shunt activity in our experiment is minimal.

## Acknowledgements

Ngala kaaditj Whadjuk Nyungar moort, kura yeye, keyen kaadak nidja boodja (*We acknowledge the Whadjuk Nyungar people, past and present, as the original custodians of the land where we live and work).* AB was sponsored by the Faculty of Agriculture, University of Tabriz, Iran, to visit the University of Western Australia.

## Author Contributions

A.B. and N.L.T. conceived the research plans; A.B. performed the experiments; A.B. and N.L.T. wrote the article.

